# A counterselectable sucrose sensitivity marker permits efficient and flexible mutagenesis in *Streptococcus agalactiae*

**DOI:** 10.1101/499822

**Authors:** Thomas A. Hooven, Maryam Bonakdar, Anna B. Chamby, Adam J. Ratner

## Abstract

*Streptococcus agalactiae* (group B *Streptococcus;* GBS) is a cause of severe infections, particularly during the newborn period. While methods exist for generating chromosomal mutations in GBS, they are cumbersome and inefficient and present significant challenges if the goal is to study subtle mutations such as single base pair polymorphisms. To address this problem, we have developed an efficient and flexible GBS mutagenesis protocol based on sucrose counterselection against levansucrase (SacB) expressed from a temperature-selective shuttle vector. GBS containing the SacB expression cassette demonstrate lethal sensitivity to supplemental sucrose whether the plasmid DNA is replicating outside of the chromosome or has been integrated during a crossover event. Transmission electron microscopy shows that SacB-mediated lethal sucrose sensitivity results from accumulation of inclusion bodies that eventually lead to complete degradation of normal cellular architecture and subsequent lysis. We used this new mutagenesis technique to generate an in-frame, allelic exchange knockout of the GBS sortase gene *srtA,* demonstrating that >99% of colonies that emerge from our protocol had the expected knockout phenotype and that among a subset tested by sequencing, 100% had the correct genotype. We also generated barcoded nonsense mutations in the *cylE* gene in two GBS strains, showing that the approach can be used to make small, precise chromosomal mutations.

**Importance:** The ability to generate chromosomal mutations is fundamental to microbiology. Historically, however, GBS pathogenesis research has been made challenging by the relative genetic intractability of the organism. Generating a single knockout in GBS using traditional techniques can take many months, with highly variable success rates. Furthermore, traditional methods do not offer a straightforward way to generate single base pair polymorphisms or other subtle changes, especially to noncoding regions of the chromosome. We have developed a new sucrose counterselection-based method that permits rapid, efficient, and flexible GBS mutagenesis. Our technique requires no additional equipment beyond what is needed for traditional approaches. We believe that it will catalyze rapid advances in GBS genetics research by significantly easing the path to generating mutants.

## Introduction

*Streptococcus agalactiae* (group B *Streptococcus;* GBS) is the most common cause of neonatal sepsis and meningitis (1–3). It can also cause serious infections in adults (4, 5) and in several animal species, including fish, which can be a source of zoonotic transmission (6–9).

GBS is not naturally competent under laboratory conditions (10) and exhibits low rates of spontaneous genetic recombination (11). Genetic studies of GBS have mostly relied on generation of allelic exchange knockouts using mutagenesis cassettes cloned into a temperature-sensitive shuttle vector (12–14). The mutagenesis cassette typically consists of an antibiotic resistance marker with upstream and downstream homology arms matching the chromosomal regions adjacent to the target gene. A second antibiotic resistance marker on the plasmid, outside of the mutagenesis cassette, confers dual resistance to transformed cells.

After electroporation and transformation of competent GBS with the mutagenesis vector, transformed clones are initially grown at a permissive temperature, which allows extrachromosomal replication of the plasmid. A subsequent shift to a higher, non-permissive temperature selects against extrachromosomal plasmid replication, leaving only cells where a crossover event at one of the homology arms has resulted in plasmid integration into the chromosome.

In order to achieve allelic exchange, a second crossover event must then occur at the other homology arm, followed by plasmid expulsion from the cell. Successful completion of these steps is detected by screening individual colonies for a specific antibiotic resistance phenotype: retained antibiotic resistance from the mutagenesis cassette marker with sensitivity to the second antibiotic due to loss of the plasmid during growth at the non-permissive temperature (12).

Without effective counterselection against the plasmid, however, detection of the second crossover event—a stochastic and often rare occurrence—is inefficient. Particularly if the desired mutant has a fitness disadvantage compared to wild type, identification of an allelic exchange mutant may require manual screening of hundreds or thousands of individual colonies (15).

Furthermore, the existing approach limits the types of mutants that can be generated. Since the final screen depends on persistence of one antibiotic resistance phenotype with loss of a second, mutants produced using traditional techniques must include an antibiotic resistance marker on the chromosome. Small-scale and unmarked alterations, such as single nucleotide polymorphisms (SNP) or subtle mutations to noncoding regions, are very difficult to obtain.

Levansucrase (sucrose: 2,6-β-D-fructan 2,6-β-D-fructosyltransferase) is an enzyme present in multiple bacterial species and has been extensively studied in *Bacillus subtilis* (16–21). Encoded by the *sacB* gene, secreted *B. subtilis* levansucrase polymerizes sucrose into the branched fructan polymer known as levan (16, 19). While the exact function of levansucrase in *B. subtilis* is unknown, the levan that it generates is believed to serve a structural or nutrient role for the cell (17).

In other bacterial species, expression of *B. subtilis sacB* confers lethal sensitivity to sucrose (22, 23). The mechanism of sucrose toxicity is believed to be from either intracellular or extracellular accumulation of levan, with resultant disruption of normal cellular processes. Sucrose sensitivity from *sacB* expression has been used as counterselection in conjunction with plasmid-based mutagenesis systems to isolate mutants in several bacterial species (22, 23). To date, however, the technique has not been described in GBS.

Here we report development and validation of a flexible and efficient counterselection system to make targeted mutations in GBS using a sacB-containing, temperature sensitive, broad host range plasmid. A schematic of this new system is presented in **Figure 1**. We show that the system can be used to generate marked and unmarked mutations in multiple GBS strains. In our experience, use of this technique dramatically decreases the labor and time required to generate mutants, allowing accelerated discovery.

**Figure 1.**
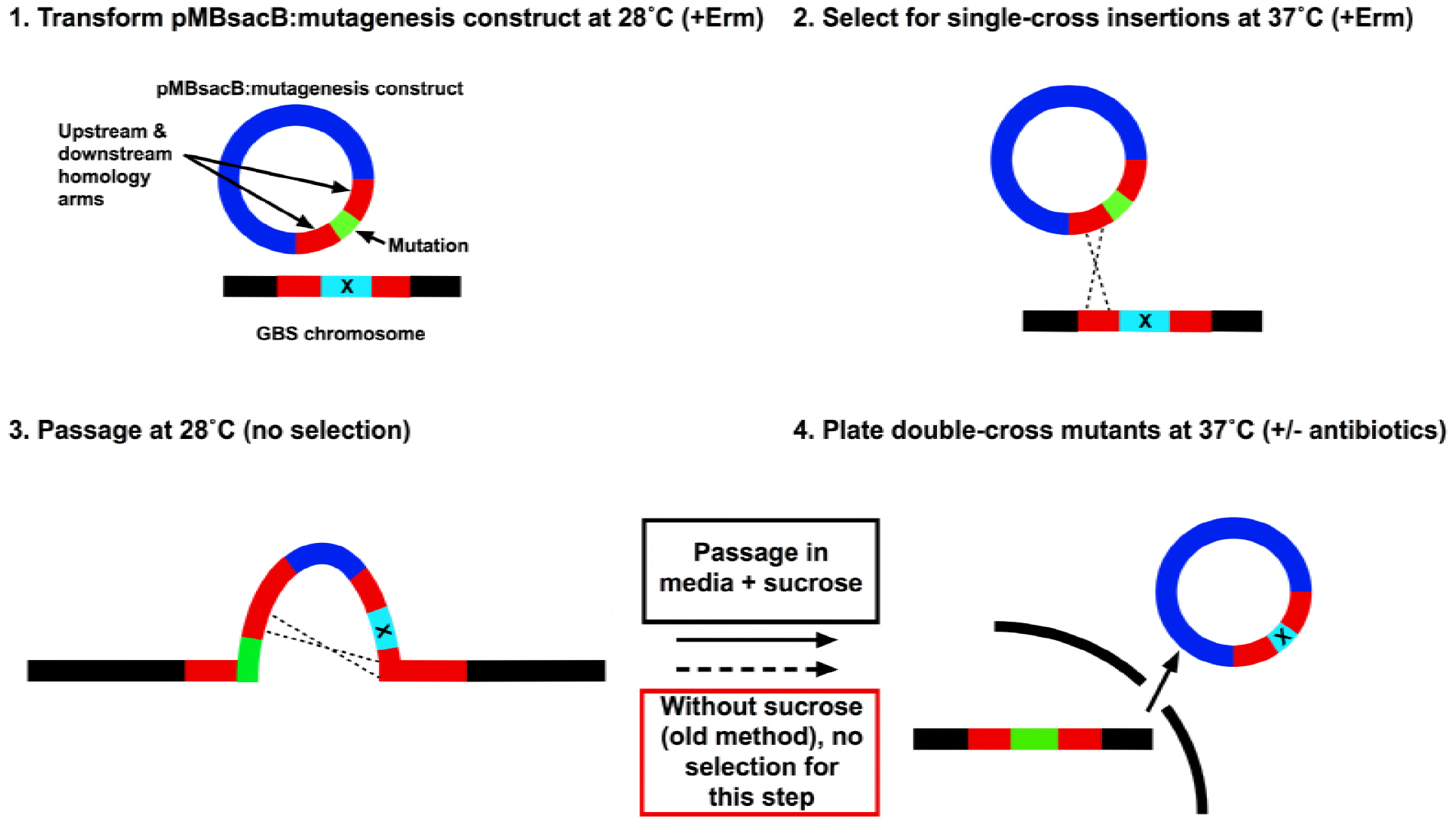
Schematic of pMBsacB-mediated GBS mutagenesis: Following insertion of a mutagenesis cassette into pMBsacB and transformation of GBS at 28 °C (1), the transformed strain is used to seed a 37 °C culture, selecting for single-crossover events at one of the homology arms (2). After removal of antibiotic selection and growth at 28 °C to promote a second crossover event and plasmid expulsion (3), sucrose is added as counterselection to isolate the desired allelic exchange clones (4). In the absence of sucrose counterselection, step 4 resembles traditional mutagenesis techniques (red box), in which identification of the allelic exchange depends on chance discovery of a clone with spontaneous loss of plasmid-based antibiotic resistance.

## Results

### Construction of pMBsacB, a temperature-sensitive, sucrose counterselectable mutagenesis shuttle vector

Plasmid pMBsacB is derived from pHY304, a widely used mutagenesis shuttle vector with the temperature-sensitive broad host range origin of replication from pWV01 (12, 24).

The *B. subtilis sacB* coding sequence, complete with signal peptide sequence, was amplified from strain 168 purified genomic DNA and cloned into plasmid pOri23, which placed the *sacB* gene downstream of the p23 promoter. This plasmid is designated pSacB23, and was used in initial experiments to test *sacB* functionality in GBS (see next section).

We subsequently generated pMBsacB by amplifying the p23 promoter and *sacB* coding sequence as a single expression cassette then subcloning it into pHY304. Figure 2 shows the steps involved in developing pMBsacB.

**Figure 2.**
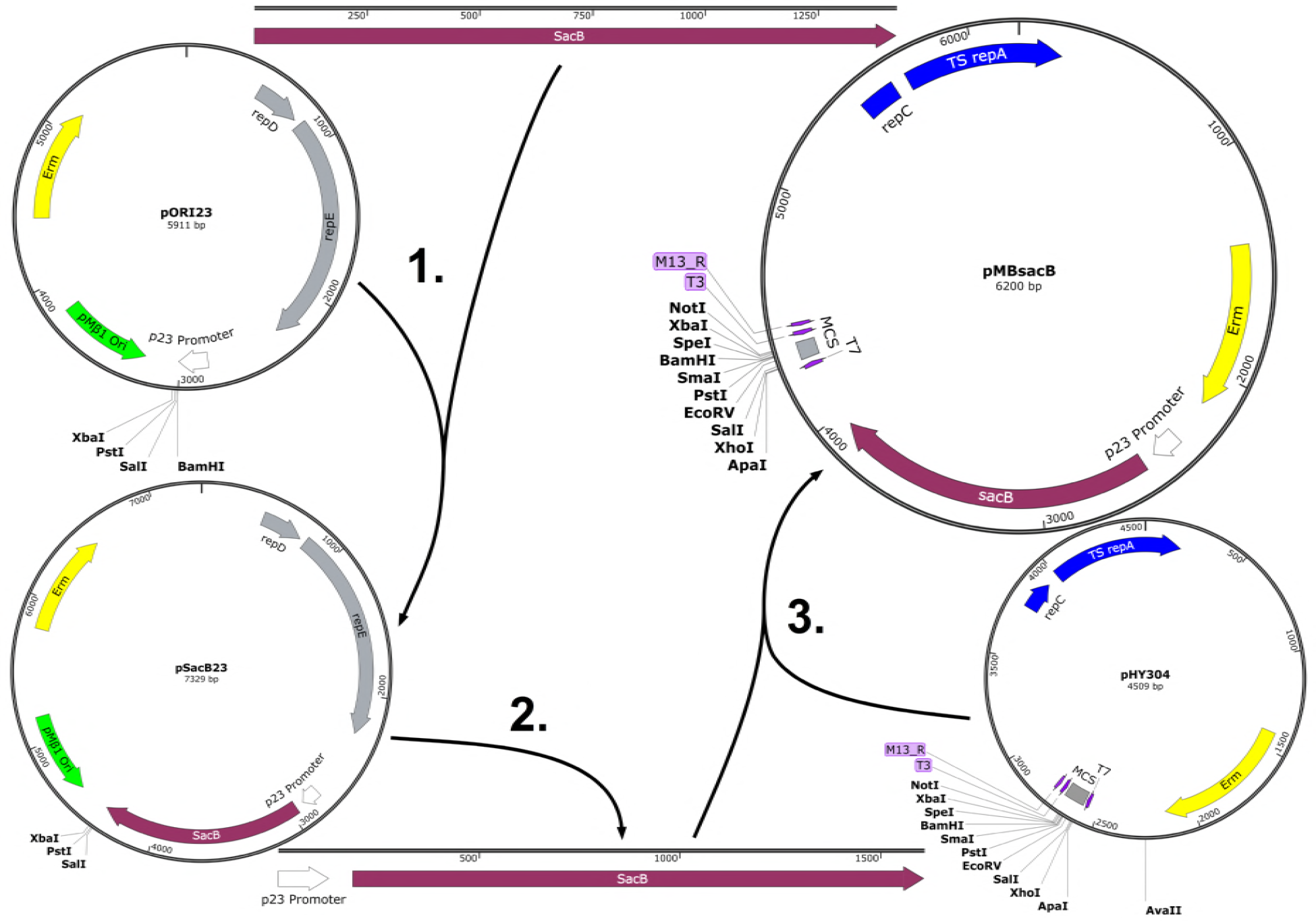
Development of pMBsacB: The *sacB* coding sequence was amplified from *B. subtilis* 168 and inserted at the *BamHI* site in pORI23, placing the gene adjacent to the p23 promoter in pSacB23 (1). The promoter-gene cassette was then amplified from pSacB23 (2) and cloned into the AvaII site of the broad host range, temperature-sensitive plasmid pHY304 (3), generating pMBsacB.

### GBS transformed with sacB-bearing plasmids show lethal sucrose sensitivity

Before transforming GBS with sacB-bearing plasmids pSacB23 and pMBsacB, we developed an electroporation protocol that did not include the use of sucrose, which is typically used as an osmoprotectant to prevent bacterial death during transformation (24, 25). We had observed very low rates of successful transformation during early trials with sucrose osmoprotection (data not shown), presumably due to sucrose-mediated toxicity.

We initially tried replacing sucrose with maltose, a structurally related disaccharide that we hoped would not be lethal to cells transformed with *sacB* plasmids. However, there was no significant increase in transformation efficiency with maltose osmoprotection (data not shown), which we attributed to *sacB* nonspecific reactivity with maltose, likely generating maltosylfructose (26, 27).

Our transformation efficiency returned to expected levels (10^-4^-10^-5^ per μg plasmid DNA for GBS strain CNCTC 10/84) with replacement of sucrose osmoprotectant with 25% (mass:mass) polyethylene-glycol (average MW 6000 daltons; PEG-6000), which we dissolved in rich transformation media for competent cell outgrowth and in the wash solution in which we store and electroporate competent GBS.

GBS strain 10/84 transformed with pSacB23 or pMBsacB showed significant growth defects on solid media with supplemental 0.75 M sucrose (**Figure 3A**). We also tested the sensitivity of planktonic 10/84:pMBsacB to sucrose added to liquid media, and found significant growth impairment. Planktonic and solid media exposure could also be combined (**Figure 3B**). 10/84 transformed with pHY304, by contrast, did not demonstrate significant sucrose sensitivity.

**Figure 3.**
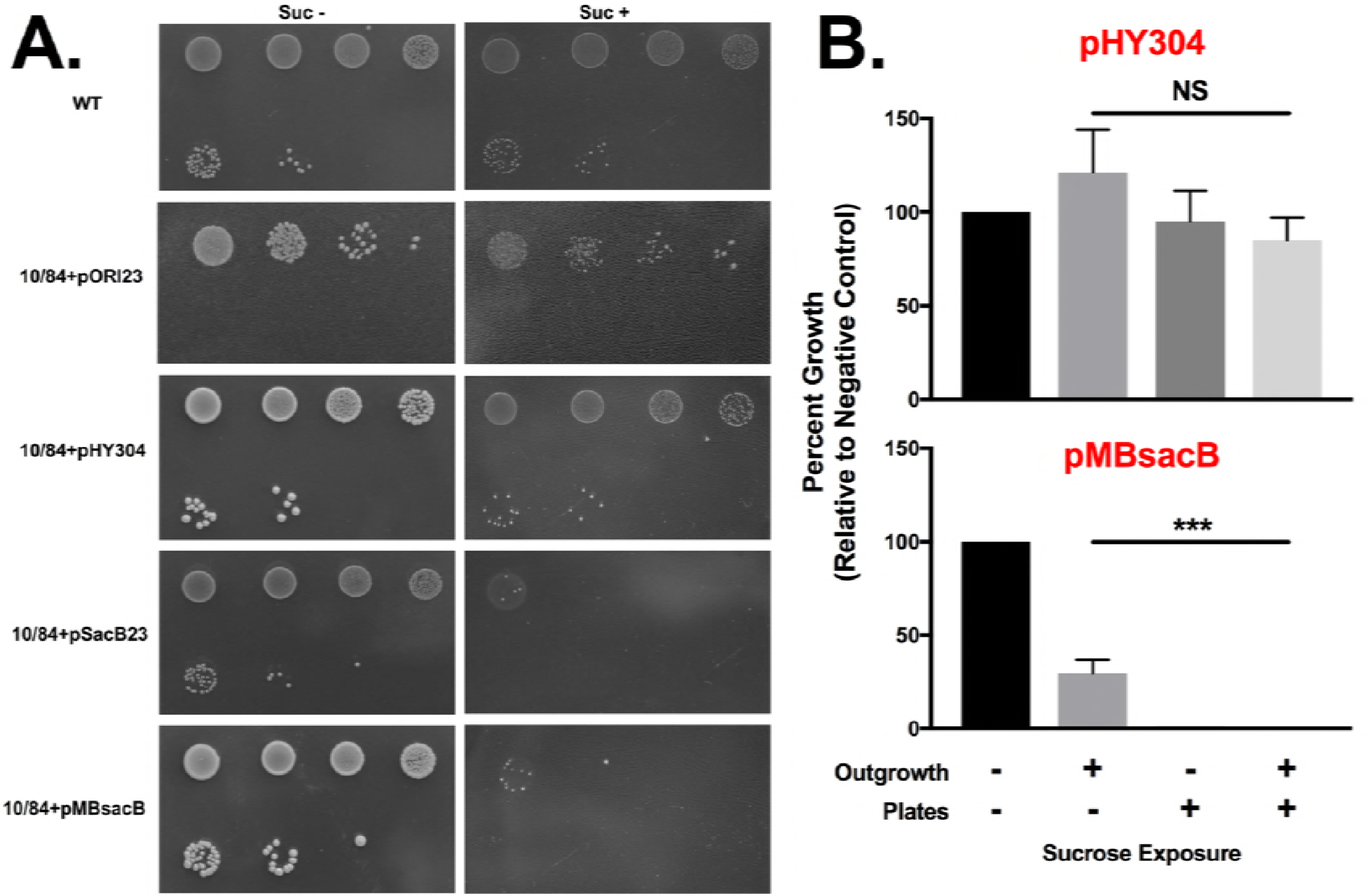
*sacB* confers lethal sucrose sensitivity in GBS: Stationary phase WT GBS 10/84 or 10/84 transformed with plasmids used in this study was serially 10-fold diluted and plated on TS agar plates with appropriate antibiotic selection with or without supplemental 0.75 M sucrose (A). The first dilution shown is 10^-1^. pMBsacB-based sucrose sensitivity functions in liquid culture and on solid media, whereas pHY304 does not affect GBS survival in sucrose in either growth condition (B; *** p < 0.0001, ANOVA).

In order to directly visualize the phenotypic effect of sucrose exposure on GBS expressing SacB, we performed transmission electron microscopy on 10/84 transformed with pMBsacB or pHY304 and exposed to sucrose or control conditions. As shown in **Figure 4A-C**, sucrose exposure of 10/84 with pMBsacB resulted in abnormal cellular morphology, with apparent intracellular accumulation of inclusion bodies, eventually resulting in complete degradation of normal cellular architecture and eventual lysis. Apart from expected cell shrinkage from sucrose-mediated osmosis, GBS transformed with the control plasmid pHY304 showed normal architecture regardless of sucrose exposure (**Figures 4D-E**). SacB expression in the absence of sucrose had no apparent effect on morphology (**Figure 4F**).

**Figure 4.**
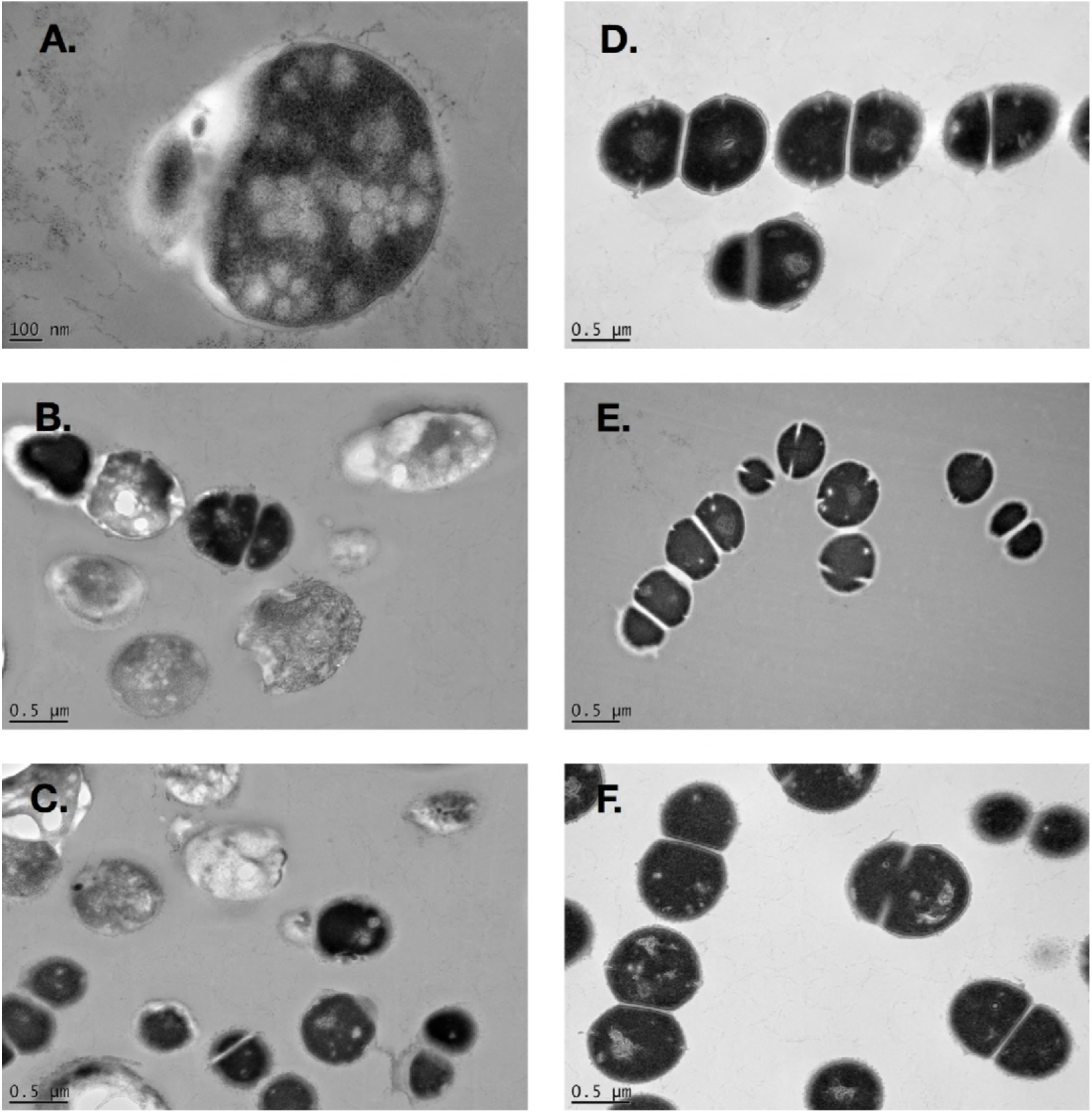
Transmission electron microscopy reveals that SacB expression under sucrose counterselection results in destructive intracellular inclusions: Early-log phase GBS strain 10/84 transformed with pMBsacB and exposed to 0.75 M sucrose for 3 hours shows intracellular inclusion bodies that lead to degraded architecture and eventual lysis (A-C). GBS transformed with the control plasmid pHY304 shows normal architecture under baseline growth conditions (D) and only expected osmotic effects when grown with supplemental sucrose (E). SacB expression in the absence of sucrose counterselection has no apparent effect on GBS morphology (F).

### Sucrose counterselection improves the efficiency of allelic exchange mutagenesis

To test whether sucrose counterselection could be used in GBS to produce allelic exchange mutants, we applied our system to deleting a gene that we and others had experience knocking out: the sortase gene *srtA* (28). This provided a benchmark against which efficiency of counterselection-assisted mutagenesis could be measured.

We used overlap extension PCR to generate a *srtA* mutagenesis cassette *(dSrtA)* in which approximately 800-bp homology arms flank an in-frame chloramphenicol acetyltransferase gene (cat), which confers chloramphenicol resistance, replacing the *srtA* coding sequence. This cassette was subcloned into pHY304 and pMBsacB, then used for transformation of 10/84 using PEG-6000 osmoprotection.

After transformation with either pHY304:dSrtA or pMBsacB:dSrtA, single-cross intermediate strains were produced by transitioning liquid cultures from 28 °C to 37 °C in the presence of erythromycin selection. Once chromosomal insertion was confirmed by PCR, the pHY304 and pMBsacB single-cross strains were serially passaged at 28 °C with no antibiotics or counterselction. At that point, each of the two cultures were used to seed two new cultures with chloramphenicol at 37 °C, one containing 0.75 M sucrose and the other a non-sucrose control.

Each of the four cultures (pHY304:dSrtA and pMBsacB:dSrtA, with and without sucrose) was passaged three times at 37 °C with chloramphenicol selection. Serial dilutions were then plated on chloramphenicol-containing solid media with or without erythromycin selection. While the overall CFU concentration did not differ significantly between conditions—as indicated by equal growth on non-selective media—there was a dramatic difference between pMBsacB grown in the presence of sucrose and the other three outgrowth conditions. Exposure of the pMBsacB single-cross strain to sucrose eliminated virtually all erythromycin-resistant survival, suggesting successful counterselection (**Figure 5A-B**). PCR of the *cat* gene generated the expected 660-bp gel electrophoresis bands when genomic DNA from colonies that survived counterselection was used as template, but not wild type GBS DNA (**Figure 5C**). Sanger sequencing of DNA amplified using PCR primers that bind outside of the *dSrtA* homology arms confirmed that the *srtA* gene had been replaced by *cat,* as intended (**Figure 5D**). The pHY304 single-cross strain was not responsive to sucrose, and the pMBsacB erythromycin-resistant CFUs survived well in the absence of sucrose exposure.

**Figure 5.**
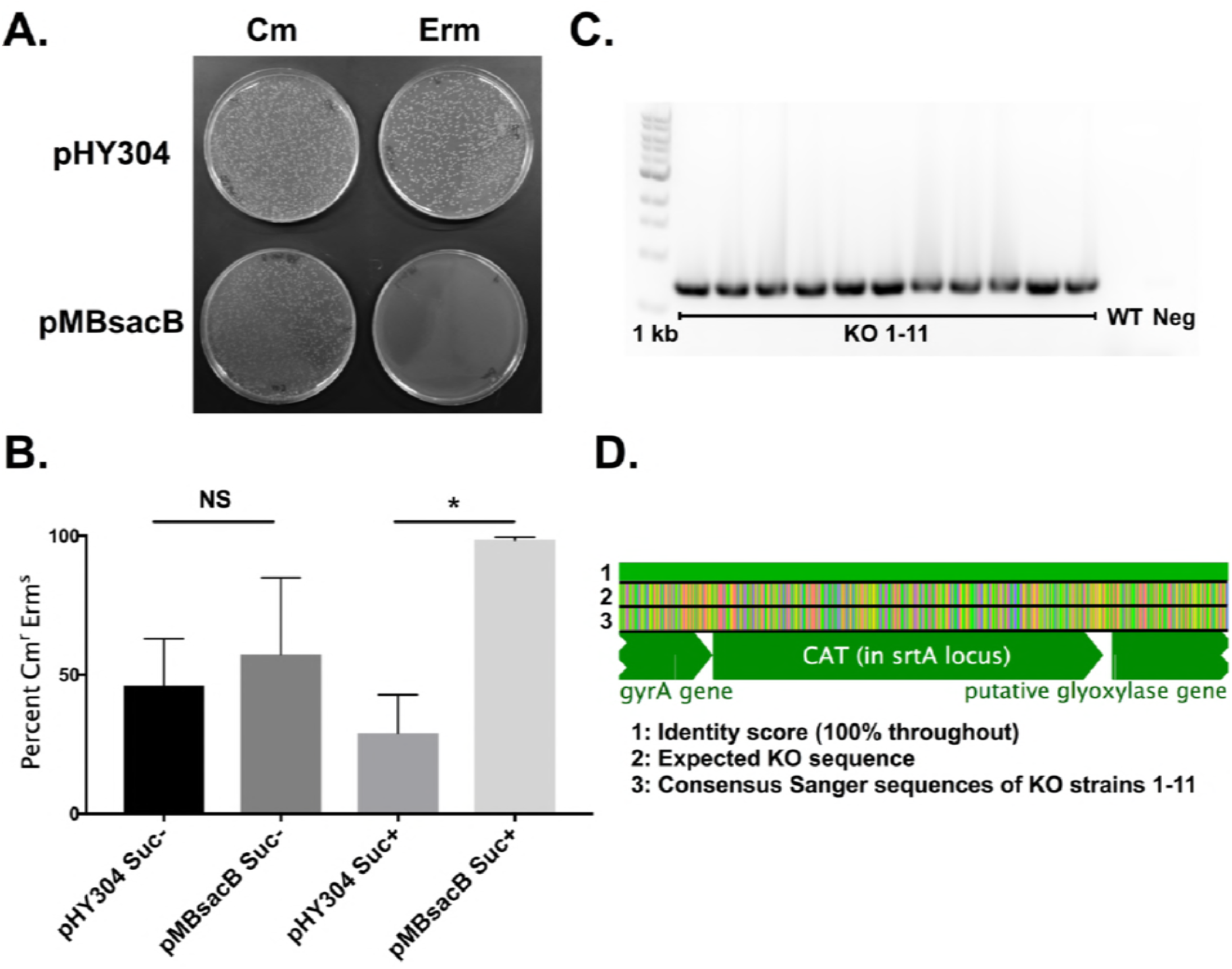
Allelic exchange mutagenesis of the *srtA* gene is made more efficient by using pMBsacB: Sucrose exposure of single-cross pMBsacB:*dsrtA* knockout intermediates results in selection against erythromycin-resistant clones, which is not seen with the pHY304:*dsrtA* single-cross strain (A). Recovery of phenotypically correct knockout clones is significantly more efficient using pMBsacB than when mutagenesis is performed with pHY304 (B; * p < 0.05, T test). PCR of *cat* results in successful amplification of genomic DNA from 11 knockout (KO) candidates, but not from wild type (WT) or template-negative (Neg) controls (C). Sanger sequences of the *srtA* region amplified using primers outside of the mutagenesis cassette were combined to generate a consensus sequence, which was aligned to the expected knockout template. Erm=erythromycin, Cm=chloramphenicol, suc=sucrose.

### The pMBsacB mutagenesis system permits efficient generation of unmarked mutations in multiple GBS strains

An effective counterselection system could facilitate generation of unmarked mutations, opening the possibility of performing sequential genome edits to produce complex mutants.

To explore whether pMBsacB could accelerate generation of unmarked mutations, we developed a mutagenesis cassette against the *cylE* gene, which encodes a key biosynthetic enzyme responsible for production of the GBS pigmented hemolytic toxin β-hemolysin/cytolysin (29–31). The mutagenesis cassette features a premature stop codon surrounded by a set of silent mutation SNPs, establishing a unique barcode that would not likely arise through spontaneous mutation. The barcoded stop codon was flanked by 500-bp upstream and downstream homology arms (**Figure 6A**).

**Figure 6.**
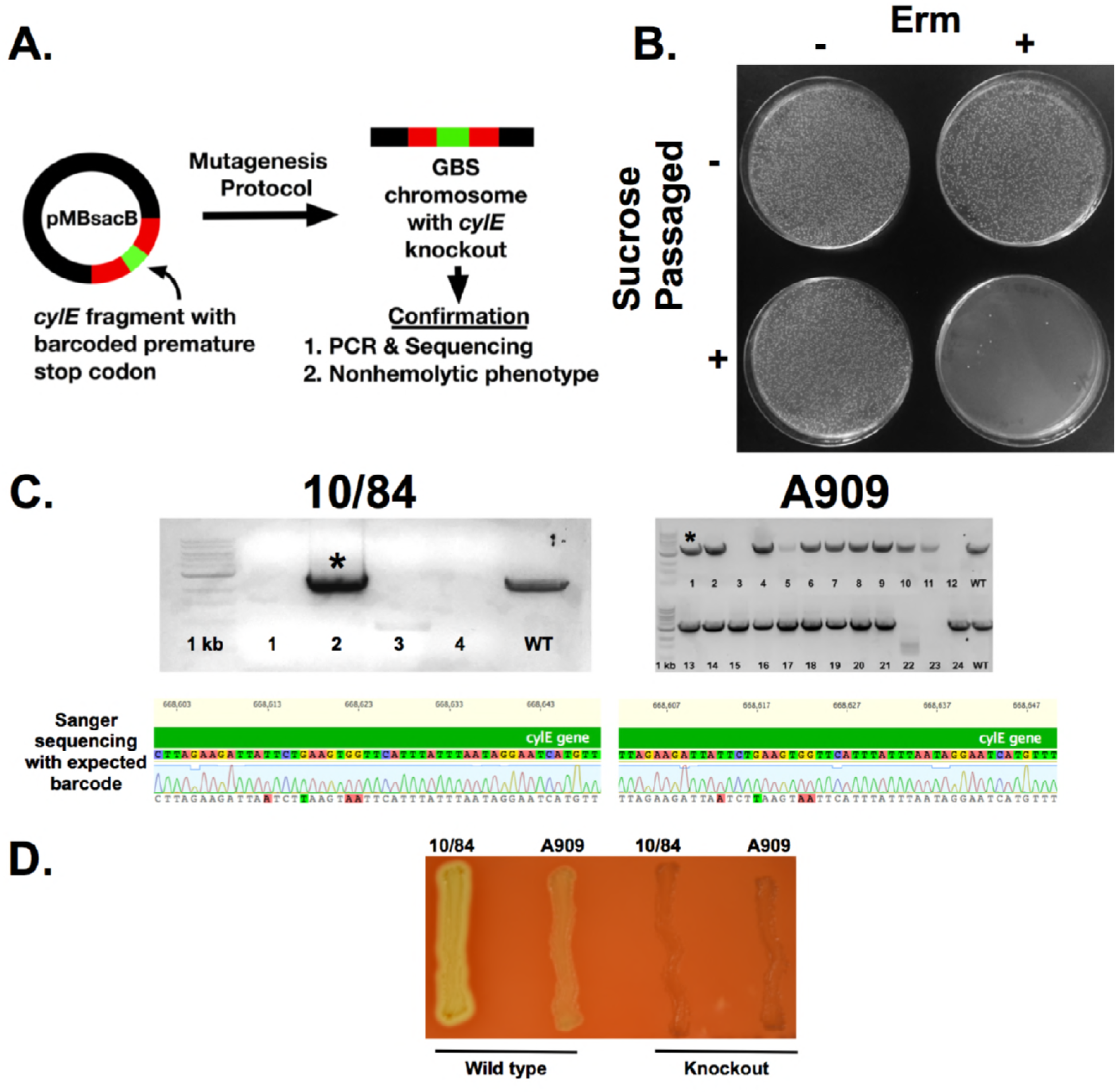
pMBsacB permits efficient generation of unmarked mutations in the *cylE* gene: After generation of a *cylE* mutagenesis cassette bearing a premature stop codon barcoded by additional silent SNPs, the cassette was cloned into pMBsacB and used to knockout β-hemolysin/cytolysin expression in GBS 10/84 and A909 (A). Sucrose counterselection resulted in near-complete elimination of erythromycin-resistant CFUs (B). Clones that survived counterselection were screened for the barcoded mutation by performing PCR with cylE_farout_F/R primers and Sanger sequencing of the amplified genomic region of interest with dCylE_Conf_F (C). Representative traces are shown for the bands marked by asterisks. The recovered knockout strains had the expected non-hemolytic phenotype when plated on 5% sheep's blood agar (D). Erm=erythromycin.

We cloned this cassette into pMBsacB, generating pMBsacB:dCylE, which was then transformed into two GBS strains, 10/84 and A909. After generation of the single-cross intermediate, which was non-hemolytic (data not shown), we used sucrose counterselection to isolate putative double-cross GBS, which could be either wild type—due to auto-excision of the plasmid at the same homology arm as crossed over during insertion—or a *cylE* knockout.

Counterselection against the single-cross strain again resulted in near-complete elimination of erythromycin resistant CFUs (**Figure 6B**).

When plated on solid media without antibiotic selection, the counterselection culture yielded an equal mix of pigmented and non-pigmented colonies, supporting the concept that the plasmid could auto-excise in one of two ways, only one of which would result in the non-pigmented phenotype expected of the knockout. If not exposed to sucrose, the single-cross intermediate retained erythromycin resistance and the colonies were uniformly nonpigmented, reflecting the fact that the population consisted almost entirely of unchanged, single-cross CFUs.

In the 10/84 experiment, we selected four non-pigmented colonies from the non-selective plate on which the sucrose-exposed culture had been grown. These were used for genomic DNA purification, followed by amplification of the *cylE* coding sequence and Sanger sequencing. One of the four had the correct barcoded sequence, whereas the other three did not properly amplify during PCR, suggesting that the plasmid auto-excised in a manner that left a partial sequence deletion (**Figure 6C**).

In the case of A909, 20 out of 24 non-pigmented isolates properly amplified by PCR. We sequenced ten of these, nine of which had the expected barcoded premature stop codon (**Figure 6C**). Both 10/84 and A909 *cylE* knockout strains generated using pMBsacB:dCylE showed the anticipated non-pigmented, non-hemolytic phenotype (**Figure 6D**).

Unintended deletions during the final plasmid-excision is a phenomenon that we have subsequently observed in other mutagenesis experiments, suggesting that all mutants generated with pMBsacB must be confirmed by some combination of PCR and sequencing to ensure the desired genotype.

## Discussion

Reliable methods for creating specific mutations are central to microbiological discovery. Existing methods for doing so in GBS have been limited in multiple respects. Without counterselection, the final screening step for plasmid auto-excision and curing is unreliable and inefficient, since it depends on random identification of a low-probability biological event. Furthermore, most current methods rely on replacement of a coding sequence with an antibiotic resistance marker; this does not support rapid generation of small changes to the chromosome, such as individual SNPs. An earlier counterselection-based approach to GBS mutagenesis was limited by the fact that it required an already mutated background strain, which is not ideal for pathogenesis work (15). Together, these barriers make isolation of complex mutants with multiple, subtle chromosomal changes infeasible.

Our *sacB*-mediated counterselection system overcomes these limitations. The method is simple and does not require any additional equipment or experience beyond what is required for traditional approaches.

We validated our system by generating two knockouts: one (*srtA*) involved allelic exchange with a chloramphenicol resistance marker, while the other *(cylE)* demonstrated the ability of our technique to produce small chromosomal changes at the single nucleotide level. In order to confirm that the system works in multiple GBS strains from different serotypes (which can show phenotypic variability under the same growth conditions), we generated the same *cylE* mutation in A909 (serotype Ia) and 10/84 (serotype V). We noted that A909 grew less robustly on 0.75 M sucrose than 10/84; so for the A909 *cylE* mutation, we used 0.5 M sucrose counterselection. When using this system in different strains, it is important to optimize the counterselection conditions prior to starting a new mutation.

As **Figure 5** shows, the *srtA* knockout can be generated at reasonable rates even without counterselection. There was considerable variability from one experimental replicate to the next, but the mean recovery rate of knockouts in the sucrose-negative conditions was 29-57%, regardless of whether the mutagenesis plasmid was pHY304 or pMBsacB. In contrast, the mean recovery rate in the pMBsacB sucrose-positive condition was 99%, with low variability.

In the case of low-fitness mutations, however, rates of recovery without counterselection can be much lower (32). The last step in the mutagenesis workflow—auto-excision and curing of the plasmid—essentially establishes a competition assay between the single-cross strain and the intended mutant (12). If the mutant has a survival defect, the odds of randomly selecting a mutant colony from among the single-cross population is very low. By shrinking the single-cross background, the pMBsacB counterselection system increases the odds of isolating the desired mutant. Particularly when the goal is mutation of high-fitness genes, we have found that it is important to confirm sucrose sensitivity of the transformant and single-cross intermediate, since spontaneous mutations in the *sacB* gene could lead to escape of unmodified high-fitness genes once sucrose counterselection is applied.

In the case of unmarked mutations, where no new antibiotic resistance is introduced into the chromosome, the bacterial population after counterselection is expected to be a mixture of the desired mutant and wild type bacteria that reverted following plasmid auto-excision. This occurred in our *cylE* knockout experiment, where roughly equal fractions of the post-counterselection population showed knockout and wild type β-hemolysin/cytolysin pigmentation phenotypes.

This means that for mutations that do not cause an easily assayable phenotype (such as pigment expression), sequence-based confirmation will be necessary. There are several possible ways to perform this confirmation. Here, we used PCR followed by Sanger sequencing, but probe-or qPCR-based SNP assays are alternative strategies. Confirmation by some means is important, given that plasmid auto-excision can leave unintended deletions, as was the case in several colonies in our *cylE* experiment.

In summary, we have presented a new, straightforward counterselection-based approach to generating flexible mutations in GBS. Our future plans for this work involve modifications to coding and non-coding sequences, including promoter alterations, addition of fluorescent and affinity tags to natively expressed genes, and generation of multiple knockout strains.

## Methods

### Bacterial strains and growth conditions

GBS strains CNCTC 10/84 (serotype V, sequence type 26) and A909 (serotype Ia, sequence type 7) and *B. subtilis* strain 168 were maintained as frozen glycerol stocks and were grown on tryptic soy (TS; Fisher Scientific product number DF0370-17-3) agar plates or stationary in TS broth at 37 °C or 28 °C. Chemically competent *Escherichia coli* DH5a were purchased from New England Biolabs (product number C2987H) and were stored and transformed according to manufacturer instructions. *E. coli* growth was at 37 °C unless transformed with a temperature-sensitive plasmid, in which case the growth temperature was 28 °C. Antibiotic concentrations were as follows: erythromycin 5 μg/mL (GBS) or 300 μg/mL *(E. coli),* chloramphenicol 1 μg/mL (GBS) or 10 μg/mL *(E. coli).* 0.5 M and 0.75 M sucrose-containing broth and solid media were prepared by diluting a filter-sterilized 2 M sucrose stock solution in appropriate media and adding any necessary antibiotics to the final mixture.

### Cloning technique

All shuttle vectors and derivatives used in this study were initially cloned into *E. coli* DH5α, from which plasmid DNA for downstream applications was purified using the Qiagen QIAprep Miniprep kit (product number 27104) according to manufacturer instructions.

### Isolation of genomic DNA

Genomic DNA from GBS and *B. subtilis* was isolated using the Applied Biosystems MagMAX CORE kit (product number A32700) with a KingFisher magnetic bead processing system according to manufacturer instructions, with the following minor modifications. Overnight liquid culture volumes were 1-10 mL. After pelleting by centrifugation at 3200 x g, the bacteria were lysed in a solution containing 100 μL manufacturer-supplied proteinase K and PK buffer, 50 μL lysozyme (100 mg/mL in water), and 5 μL mutanolysin (10 kU in 2 mL 0.1 M potassium phosphate buffer pH 6.2). Lysis was performed at 37 °C for 30 minutes, then 55 °C for 30 minutes, followed by a 2-minute centrifugation at 3200 x g. The rest of the extraction followed manufacturer instructions, using the MagMAX CORE Flex KingFisher protocol file.

### Construction of pSacB23 and pMBsacB

The *sacB* coding sequence was amplified from *B. subtilis* 168 genomic DNA using primers sacB_pORI23_F and sacB_pORI23_R, which contain Gibson assembly overhang sequences compatible with pORI23 digested with *BamHI.* Successful clones were identified by Sanger sequencing (data not shown), after which the p23 promoter-sacB cassette was amplified with p23sacB_cassette_F and p23sacB_cassette_R and subloned into pHY304 digested with *AvaII* (see **Figure 1**). Outgrowth of the pMBsacB cloning reaction was performed at 28 °C and successful clones were identified with Sanger sequencing (data not shown).

### Transformation of GBS with pSacB23 and pMBsacB

Electrocompetent GBS suitable for transformation with sacB-containing plasmids were prepared following methods outlined by Holo and Nes (25) and Framson et al. (24) modified to prevent toxicity from sucrose osmoprotectant.

Single 10/84 or A909 colonies from TS agar plates were used to seed 5 mL M17 (BD Difco 218561) + 0.5% glucose liquid cultures, which were grown at 37 °C to stationary phase. 500 μL from these cultures were then used to seed filter sterilized 50 mL M17 + 0.5% glucose, 2.5% (A909) or 0.6% (10/84) glycine, and 25% (mass:mass) PEG-6000.

Following overnight growth at 37 °C, this culture was diluted in pre-warmed 130 mL of the same media and allowed to grow for 1 hr. The entire volume was pelleted at 3200 x g, then washed twice in ice cold 25% PEG-6000 + 10% glycerol. Following these washes, the samples were resuspended in 1 mL of wash solution and either used immediately for transformation or stored in aliquots at −80 °C.

Electroporation and transformation of competent GBS was performed as described by Holo and Nes (25), except that sucrose in the outgrowth media was replaced with 25% PEG-6000, and— in the case of pMBsacB—outgrowth was performed at 28 °C instead of 37 °C.

### Sucrose killing assays

Wild type GBS or transformants were grown overnight in appropriate antibiotic selection without supplemental sucrose. For killing assays on agar plates, overnight cultures were serially diluted and plated directly on TS agar with appropriate antibiotic selection, with or without supplemental sucrose. CFU quantification was performed after 1-2 days of growth. For killing assays during planktonic growth, the overnight cultures were diluted 1:50 in broth without sucrose, grown to log phase, then exposed to sucrose supplementation (or control outgrowth with only sterile water added to the broth) for two hours, at which time the cultures were diluted and plated on appropriate solid media for CFU quantification after 1-2 days of growth.

### Transmission electron microscopy

10/84 transformed with pMBsacB or pHY304 was grown to mid-log phase in selective broth. That culture was used to seed a new culture with or without supplemental sucrose. After outgrowth to early-mid log phase, the bacteria were fixed with 2.5% glutaraldehyde and 2% paraformaldehyde and washed with cacodylate buffer (50 mM, pH 7.2), then post fixed with 2% osmium tetroxide. Bacteria were embedded in 2% agar, then cut and stained in the dark with 0. 5% (w/v) uranyl acetate. Samples were dehydrated with alcohol, transferred to propylene oxide/Epon mixtures and finally embedded in EMbed 812 (Electron Microscopy Sciences, Hatfield, PA). Thin sections were cut, adsorbed on electron microscope grids, and stained with uranyl acetate and lead citrate. Stained grids were then imaged in a Philips CM12 electron microscope (FEI; Eindhoven, The Netherlands) and photographed with a Gatan (4kx2.7k) digital camera (Gatan Inc., Pleasanton, CA).

### Construction of pMBsacB:dSrtA and pMBsacB:dCylE

The *dSrtA* mutagenesis cassette consisted of the *cat* gene with 822-bp upstream and 795-bp downstream homology arms that matched the 10/84 chromosomal regions flanking the *srtA* gene. The three fragments were amplified from template DNA (10/84 genomic DNA or pDC123, a shuttle vector containing *cat).* The primers used to generate the upstream, coding, and downstream cassette regions of the mutagenesis cassette were dSrtA_US_F/R, cat_F/R, and dSrtA_DS_F/R, respectively.

Next, overlap extension PCR was used to join the three regions (28). The upstream homology arm was first joined to *cat,* after which the downstream homology arm was attached. The two-fragment intermediate and the final cassette were gel extracted and used for TOPO cloning into pCR2.1-TOPO, followed by Sanger sequence confirmation (data not shown). After successful construction, the complete cassette was digested out of pCR2.1-TOPO with *NotI* and *KpnI,* then ligated with T4 ligase overnight at 16 °C into pHY304 double-digested with the same restriction enzymes. For insertion of the *dSrtA* cassette into pMBsacB, the construct was amplified out of pHY304:dSrtA using primers dSrtA_GA_F and dSrtA_GA_R, then cloned into pMBsacB at the *NotI* and *XhoI* sites using Gibson assembly.

The 1500-bp *dCylE* cassette was amplified out of GBS strain AR1598, a derivative of 10/84 in which the *cylE* gene has a barcoded premature stop codon (**Figure 6B**), using primers dCylE_GA_F and dCylE_GA_R. This amplicon was similarly cloned into the *NotI* and *XhoI* sites of pMBsacB using Gibson assembly.

### Allelic exchange using sucrose counterselection against pMBsacB

After transformation of GBS, using the method described above, with pMBsacB (or pHY304 control) bearing a mutagenesis cassette (dSrtA or *dCylE),* successful transformants were grown in TS broth with appropriate antibiotic selection (erythromycin with or without chloramphenicol) at 28 °C. Sucrose sensitivity of the transformants was confirmed by plating serial dilutions on TS agar with 0.5 M (for A909) or 0.75 M (for 10/84) sucrose and appropriate antibiotic selection at 28 °C.

To generate single-cross intermediates, transformants were serially passaged three times at 28 °C with erythromycin selection. The third passage was then used to seed another culture at 37 °C with erythromycin selection, which was grown overnight. Serial dilutions of the final culture were plated on TS agar with erythromycin at 37 °C. Individual colonies were grown and tested for sucrose sensitivity. Genomic DNA was also extracted and tested by PCR for proper vector insertion using either cylE_farout_F or srtA_farout_F, which match chromosomal sites outside of the homology arms, and pMBsacB_MCS_F, which binds pMBsacB and pHY304 upstream of the cloning sites used in this study (data not shown).

Single-cross intermediate strains with the correct sucrose sensitivity phenotype and PCR-confirmed genotype were then grown in TS broth without antibiotics at 28 °C and passaged three times in order to enrich for spontaneous double-cross events. To counterselect against pMBsacB, the third passage was used to seed TS broth with sucrose at 37 °C. In *srtA* knockout experiments, chloramphenicol was added to the sucrose-containing broth. The sucrose culture was passaged three times at 37 °C, and then serial dilutions were plated on TS agar with or without chloramphenicol. Simultaneous plating on erythromycin-containing TS agar (with or without chloramphenicol) was used to quantify the effectiveness of counterselection against pMBsacB.

Knockout candidates from the non-erythromycin plate were confirmed to be erythromycin sensitive by patching to a new plate. Genomic DNA extraction followed by PCR and Sanger sequencing confirmed plasmid excision and the correct knockout DNA sequence. For the *srtA* knockout, the region was amplified using primers srtA_farout_F/R and these amplicons were sequenced using srtA_farout_F and cat_F primers. For the *cylE* knockout, the region was amplified using cylE_farout_F/R and the barcoded mutation was confirmed by sequencing with the dCylE_conf_F primer.

pHY304 knockout controls were subjected to the same steps, including sucrose exposure, unless otherwise noted. Non-sucrose exposure control conditions were identical except that the three final passages of the single-cross strain at 37 °C were performed in the absence of sucrose.

**Table 1:**
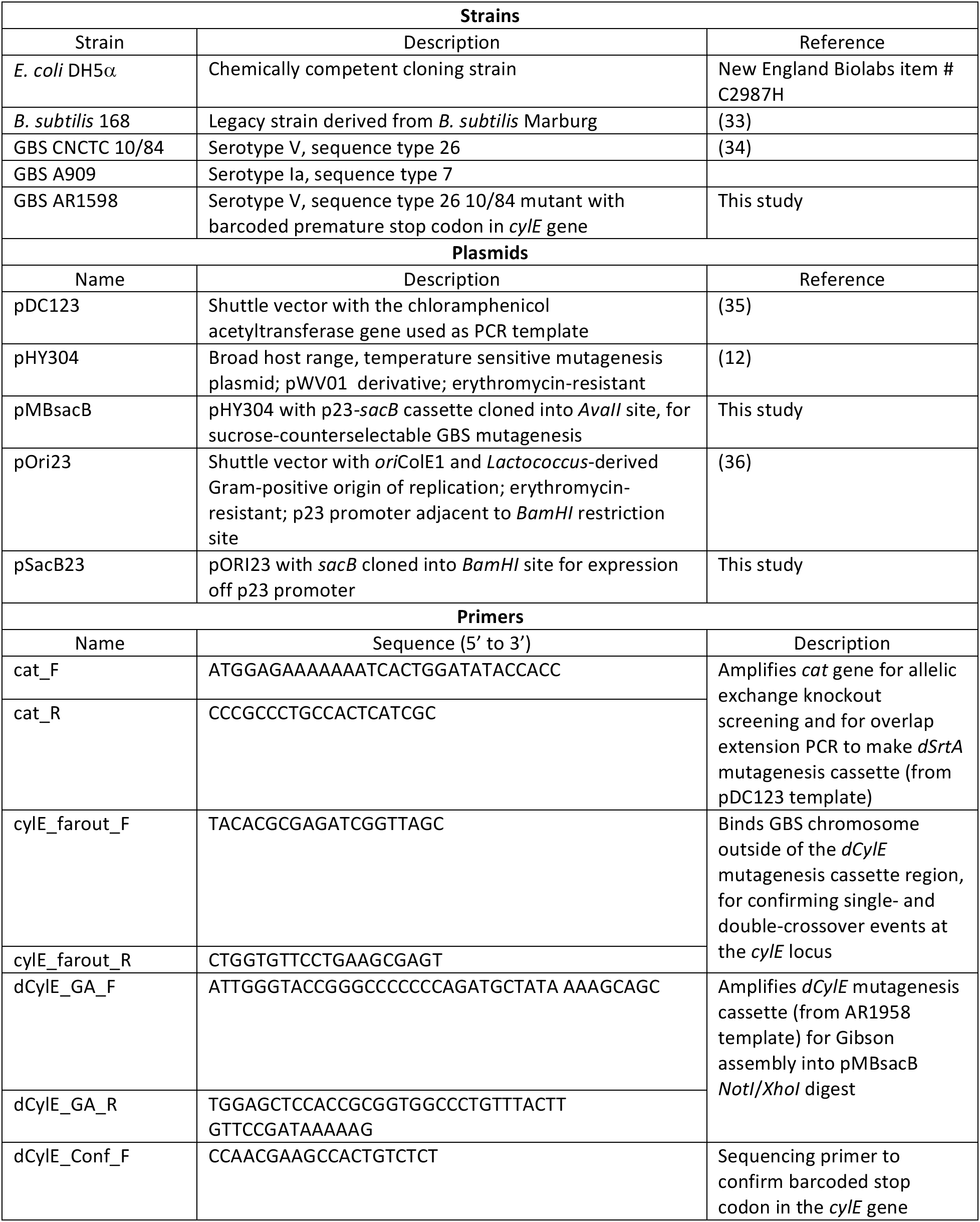
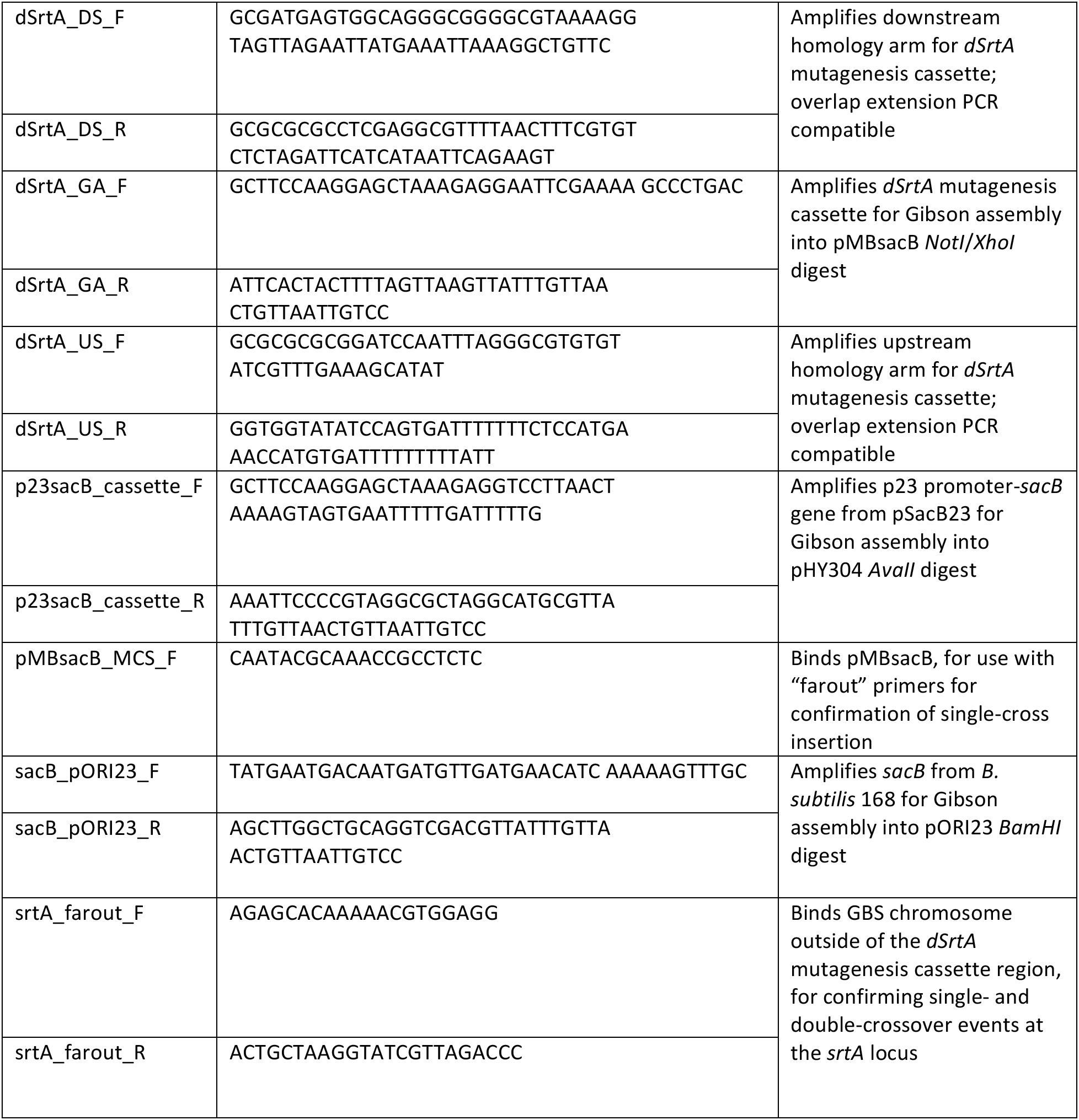
Strains, plasmids, and primers used in this study

## Acknowledgements

We thank NYU Langone Health DART Microscopy Laboratory for the consultation and assistance with transmission electron microscopy work.

## References

1. Baker CJ. 2013. The spectrum of perinatal group B streptococcal disease. Vaccine 31 Suppl 4:D3–6.

2. Stoll BJ, Hansen NI, Sánchez PJ, Faix RG, Poindexter BB, Van Meurs KP, Bizzarro MJ, Goldberg RN, Frantz ID, Hale EC, Shankaran S, Kennedy K, Carlo WA, Watterberg KL, Bell EF, Walsh MC, Schibler K, Laptook AR, Shane AL, Schrag SJ, Das A, Higgins RD, Eunice Kennedy Shriver National Institute of Child Health and Human Development Neonatal Research Network. 2011. Early onset neonatal sepsis: the burden of group B Streptococcal and E. coli disease continues. Pediatrics 127:817–826.

3. Schrag SJ, Farley MM, Petit S, Reingold A, Weston EJ, Pondo T, Hudson Jain J, Lynfield R. 2016. Epidemiology of Invasive Early-Onset Neonatal Sepsis, 2005 to 2014. Pediatrics 138:e20162013–e20162013.

4. Muñoz P, Llancaqueo A, Rodríguez-Créixems M, Peláez T, Martin L, Bouza E. 1997. Group B streptococcus bacteremia in nonpregnant adults. Arch Intern Med 157:213–216.

5. Reingold A, Watt JP. 2015. Group B streptococcus infections of soft tissue and bone in California adults, 1995-2012. Epidemiol Infect 143:3343–3350.

6. Kayansamruaj P, Pirarat N, Hirono I, Rodkhum C. 2014. Increasing of temperature induces pathogenicity of Streptococcus agalactiae and the up-regulation of inflammatory related genes in infected Nile tilapia (Oreochromis niloticus). Vet Microbiol 172:265–271.

7. Botelho ACN, Ferreira AFM, Fracalanzza SEL, Teixeira LM, Pinto TCA. 2018. A Perspective on the Potential Zoonotic Role of Streptococcus agalactiae: Searching for a Missing Link in Alternative Transmission Routes. Front Microbiol 9:608.

8. Kalimuddin S, Chen SL, Lim CTK, Koh TH, Tan TY, Kam M, Wong CW, Mehershahi KS, Chau ML, Ng LC, Tang WY, Badaruddin H, Teo J, Apisarnthanarak A, Suwantarat N, Ip M, Holden MTG, Hsu LY, Barkham T, Singapore Group B Streptococcus Consortium. 2017. 2015 Epidemic of Severe Streptococcus agalactiae Sequence Type 283 Infections in Singapore Associated With the Consumption of Raw Freshwater Fish: A Detailed Analysis of Clinical, Epidemiological, and Bacterial Sequencing Data. Clin Infect Dis 64:S145–S152.

9. Tan S, Lin Y, Foo K, Koh HF, Tow C, Zhang Y, Ang LW, Cui L, Badaruddin H, Ooi PL, Lin RTP, Cutter J. 2016. Group B Streptococcus Serotype III Sequence Type 283 Bacteremia Associated with Consumption of Raw Fish, Singapore. Emerging Infect Dis 22:1970–1973.

10. Håvarstein LS, Hakenbeck R, Gaustad P. 1997. Natural competence in the genus Streptococcus: evidence that streptococci can change pherotype by interspecies recombinational exchanges. J Bacteriol 179:6589–6594.

11. Brochet M, Rusniok C, Couve E, Dramsi S, Poyart C, Trieu-Cuot P, Kunst F, Glaser P. 2008. Shaping a bacterial genome by large chromosomal replacements, the evolutionary history of Streptococcus agalactiae. Proc Natl Acad Sci USA 105:15961–15966.

12. Yim HH, Rubens CE. 1998. Site-specific homologous recombination mutagenesis in group B streptococci. Methods in Cell Science 20:13–20.

13. Sheen TR, Jimenez A, Wang N-Y, Banerjee A, van Sorge NM, Doran KS. 2011. Serine-rich repeat proteins and pili promote Streptococcus agalactiae colonization of the vaginal tract. Journal of Bacteriology 193:6834–6842.

14. Quach D, van Sorge NM, Kristian SA, Bryan JD, Shelver DW, Doran KS. 2009. The CiaR response regulator in group B Streptococcus promotes intracellular survival and resistance to innate immune defenses. Journal of Bacteriology 191:2023–2032.

15. Tamura GS, Bratt DS, Yim HH, Nittayajarn A. 2005. Use of glnQ as a counterselectable marker for creation of allelic exchange mutations in group B streptococci. Appl Environ Microbiol 71:587–590.

16. Meng G, Fütterer K. 2003. Structural framework of fructosyl transfer in Bacillus subtilis levansucrase. Nat Struct Biol 10:935–941.

17. Marvasi M, Visscher PT, Casillas Martinez L. 2010. Exopolymeric substances (EPS) from Bacillus subtilis: polymers and genes encoding their synthesis. FEMS Microbiol Lett 313:1–9.

18. Porras-Domínguez JR, Ávila-Fernández Á, Miranda-Molina A, Rodriguez-Alegria ME, Munguia AL. 2015. Bacillus subtilis 168 levansucrase (SacB) activity affects average levan molecular weight. Carbohydr Polym 132:338–344.

19. Ortiz-Soto ME, Rudiño-Piñera E, Rodriguez-Alegria ME, Munguia AL. 2009. Evaluation of cross-linked aggregates from purified Bacillus subtilis levansucrase mutants for transfructosylation reactions. BMC Biotechnol 9:68.

20. Abdel-Fattah AF, Mahmoud DAR, Esawy MAT. 2005. Production of levansucrase from Bacillus subtilis NRC 33a and enzymic synthesis of levan and Fructo-Oligosaccharides. Curr Microbiol 51:402–407.

21. Ortiz-Soto ME, Rivera M, Rudiño-Piñera E, Olvera C, López-Munguía A. 2008. Selected mutations in Bacillus subtilis levansucrase semi-conserved regions affecting its biochemical properties. Protein Eng Des Sel 21:589–595.

22. Marx CJ. 2008. Development of a broad-host-range sacB-based vector for unmarked allelic exchange. BMC Res Notes 1:1.

23. Li Y, Thompson CM, Lipsitch M. 2014. A modified Janus cassette (Sweet Janus) to improve allelic replacement efficiency by high-stringency negative selection in Streptococcus pneumoniae. PLoS ONE 9:e100510.

24. Framson PE, Nittayajarn A, Merry J, Youngman P, Rubens CE. 1997. New genetic techniques for group B streptococci: high-efficiency transformation, maintenance of temperature-sensitive pWV01 plasmids, and mutagenesis with Tn917. Appl Environ Microbiol 63:3539–3547.

25. Holo H, Nes IF. 1989. High-Frequency Transformation, by Electroporation, of Lactococcus-Lactis Subsp Cremoris Grown with Glycine in Osmotically Stabilized Media. Appl Environ Microbiol 55:3119–3123.

26. Canedo M, Jimenez-Estrada M, Cassani J, López-munguía A. 1999. Production of Maltosylfructose (Erlose) with Levansucrase fromBacillus Subtilis. Biocatalysis and Biotransformation 16:475–485.

27. Seibel J, Moraru R, Götze S, Buchholz K, Na’amnieh S, Pawlowski A, Hecht H-J. 2006. Synthesis of sucrose analogues and the mechanism of action of Bacillus subtilis fructosyltransferase (levansucrase). Carbohydr Res 341:2335–2349.

28. Lalioui L, Pellegrini E, Dramsi S, Baptista M, Bourgeois N, Doucet-Populaire F, Rusniok C, Zouine M, Glaser P, Kunst F, Poyart C, Trieu-Cuot P. 2005. The SrtA Sortase of Streptococcus agalactiae is required for cell wall anchoring of proteins containing the LPXTG motif, for adhesion to epithelial cells, and for colonization of the mouse intestine. Infection and Immunity 73:3342–3350.

29. Whidbey C, Harrell MI, Burnside K, Ngo L, Becraft AK, Iyer LM, Aravind L, Hitti J, Waldorf KMA, Rajagopal L. 2013. A hemolytic pigment of Group B Streptococcus allows bacterial penetration of human placenta. J Exp Med 210:1265–1281.

30. Rosa-Fraile M, Dramsi S, Spellerberg B. 2014. Group B streptococcal haemolysin and pigment, a tale of twins. FEMS Microbiol Rev 38:932–946.

31. Randis TM, Gelber SE, Hooven TA, Abellar RG, Akabas LH, Lewis EL, Walker LB, Byland LM, Nizet V, Ratner AJ. 2014. Group B Streptococcus β-hemolysin/cytolysin breaches maternal-fetal barriers to cause preterm birth and intrauterine fetal demise in vivo. J Infect Dis 210:265–273.

32. Link AJ, Jeong KJ, Georgiou G. 2007. Beyond toothpicks: new methods for isolating mutant bacteria. Nat Rev Micro 5:680–688.

33. Zeigler DR, Prágai Z, Rodriguez S, Chevreux B, Muffler A, Albert T, Bai R, Wyss M, Perkins JB. 2008. The origins of 168, W23, and other Bacillus subtilis legacy strains. Journal of Bacteriology 190:6983–6995.

34. Hooven TA, Randis TM, Daugherty SC, Narechania A, Planet PJ, Tettelin H, Ratner AJ. 2014. Complete Genome Sequence of Streptococcus agalactiae CNCTC 10/84, a Hypervirulent Sequence Type 26 Strain. Genome Announcements 2:e01338–14–e01338–14.

35. Chaffin DO, Rubens CE. 1998. Blue/white screening of recombinant plasmids in Gram-positive bacteria by interruption of alkaline phosphatase gene (phoZ) expression. Gene 219:91–99.

36. Que YA, Haefliger JA, Francioli P, Moreillon P. 2000. Expression of Staphylococcus aureus clumping factor A in Lactococcus lactis subsp cremoris using a new shuttle vector. Infection and Immunity 68:3516–3522.

